# MORC and MOM1 spatially constrain RNA Polymerase V chromatin positioning to shape DNA methylation landscapes

**DOI:** 10.64898/2026.08.30.748062

**Authors:** Xiaoqing He, Zheng Li, Yan Xue, Jiayuan Guo, Xuexia Liu, Suhua Feng, Zhenhui Zhong, Steven E. Jacobsen

**Affiliations:** Ministry of Education Key Laboratory for Bio-Resource and Eco-Environment, Chengdu Botanical Garden-Sichuan University Joint Laboratory for Ex Situ Conservation and Resource Utilization of Mountain Plants, College of Life Sciences, Sichuan University, Chengdu, China; Chengdu Botanical Garden, Chengdu, Sichuan, China; Department of Molecular, Cell and Developmental Biology, University of California, Los Angeles, CA 90095, USA; Peking University Institute of Advanced Agricultural Sciences, Shandong Laboratory of Advanced Agricultural Sciences at Weifang, Weifang, Shandong, 261000, China; Howard Hughes Medical Institute, University of California, Los Angeles, CA 90095, USA

## Abstract

Plant-specific RNA Polymerase V (Pol V) transcribes noncoding RNAs in the RNA-directed DNA methylation pathway, thereby influencing gene expression and genome stability by controlling *de novo* DNA methylation. However, the mechanisms governing precise chromatin localization and transcriptional activities of Pol V remain elusive. Here we show that Pol V localization is spatially constrained by the chromatin regulators microrchidia (MORC) and MORPHEUS’ MOLECULE 1 (MOM1). MORC and MOM1 promote Pol V occupancy at sites near active chromatin, whereas their loss leads to redistribution of Pol V into CMT3-enriched heterochromatin, accompanied by noncoding RNA transcription, small RNA production and DNA methylation. Our findings reveal a combinatorial model in which recruitment, spatial constraint and DNA methylation feedback collectively define Pol V chromatin distribution and epigenetic function.

## Introduction

DNA methylation is an essential mechanism that establishes silent chromatin states of transposable elements (TEs) and repetitive sequences to safeguard genome integrity. It is also critical for fine-tuning gene expression during plant development and environmental adaptation^1–3^. *De novo* DNA methylation is established by the RNA-directed DNA methylation (RdDM) pathway through DOMAINS REARRANGED METHYLTRANSFERASE (DRM) enzymes that initiate cytosine methylation at target loci. Methylation is subsequently perpetuated and reinforced by maintenance methyltransferases such as MET1 and CMT2/3 to preserve epigenetic states across cell divisions. Central to the RdDM pathway is RNA Polymerase V (Pol V), a plant-specific enzyme that evolved from RNA Polymerase II (Pol II) but diverged to fulfill a specialized role in the RdDM pathway^4,5^. Unlike Pol II, which primarily transcribes protein-coding mRNAs, Pol V synthesizes long noncoding RNAs (ncRNAs) at TEs and repetitive DNA^6–10^. These ncRNAs act as scaffolds to recruit siRNA-ARGONAUTE 4 (AGO4) complexes, which in turn direct DRM2 to catalyze *de novo* DNA methylation. The RdDM pathway is mainly located at TEs and repetitive sequences, enriched with dense DNA methylation and repressive histone modifications^11^. However, RdDM also operates near transcriptionally active regions, where it may modulate protein coding gene expression and chromatin accessibility of young TEs^12^. This broad transcription capacity suggests that Pol V plays a major role in maintaining epigenetic states^12,13^. Furthermore, protein structural studies revealed that Pol V exhibits unique transcriptional properties: while it shares Pol II’s fidelity, it lacks transcription factor-binding interfaces and instead engages in pervasive transcription, which raises the question of how Pol V navigates to chromatin and targets specific loci without canonical regulatory domains^4,10^.

Previous studies have identified several factors required for Pol V recruitment and activity. For example, the chromatin-remodeling complex DDR (DRD1: DEFECTIVE IN RNA-DIRECTED DNA METHYLATION 1, DMS3: DEFECTIVE IN MERISTEM SILENCING 3, and RDM1: RNA-DIRECTED DNA METHYLATION 1) and DRM2/3 facilitate Pol V association with chromatin, while SUVH2 and SUVH9 recognize methylated DNA and promote Pol V occupancy at repressive loci^12,14–16^. In addition, AGO4–siRNA complexes interact with nascent Pol V transcripts, reinforcing the targeting specificity of RdDM^17^. In contrast, the linker histone H1 restricts the spreading of Pol V into heterochromatin^18^. These findings highlight a multilayered network regulating Pol V occupancy and transcription. The MICRORCHIDIA (MORC) and MORPHEUS’ MOLECULE 1 (MOM1) are critical for reinforcing TE silencing and chromatin compaction, with MORC proteins acting as GHKL-type ATPases and MOM1 sharing homology to SWI2/SNF2 chromatin remodelers^19–24^. These factors collaborate to recruit and stabilize the RdDM machinery: MOM1 bridges MORC6 to Pol V at target loci, while MORC proteins act as molecular tethers to enhance DNA methylation^25^. Their colocalization at silent chromatin regions suggests a synergistic role of these factors in transcriptional repression of TEs and genes^25,26^. Genome-wide studies revealed that MORC and MOM1 occupy both transcriptionally active and inactive chromatin domains, whereas Pol V remains predominantly enriched at methylated heterochromatin^25,27,28^. This spatial discordance raises key questions about how MORC and MOM1 interface with Pol V to regulate RdDM. Additionally, zinc finger-mediated tethering of MOM1 or MORC1 to ectopic loci can recruit Pol V and establish *de novo* DNA methylation^29^. Yet, the mechanisms enabling MORC and MOM1 to guide Pol V activity across diverse chromatin contexts and the functional consequences of their interplay remain unresolved.

In this study, we uncovered essential roles for MORC and MOM1 in spatially regulating Pol V occupancy to ensure precision of DNA methylation. We demonstrate that MORC and MOM1 stabilize Pol V at AT-rich and hypomethylated loci, preventing its ectopic redistribution into heterochromatic regions. Loss of MORC and MOM1 leads to widespread Pol V relocation to heterochromatin, which in turn drives aberrant Pol V transcription, siRNA biogenesis, and DNA methylation. These findings establish MORC and MOM1 as essential modulators of Pol V targeting, acting together with SUVH2/9 and DNA methylation pathways.

## Results

### Chromatin factors define Pol V distribution

The activity of Pol V is subject to rigorous regulatory mechanisms to ensure its precise targeting to specific genomic regions to initiate RdDM^29^. Previous studies demonstrated that MORC and MOM1 proteins play an important role in tethering Pol V to RdDM sites^25,26,29^. To further investigate the influence of MORC and MOM1 proteins on the genome-wide distribution of Pol V, we conducted chromatin immunoprecipitation followed by DNA sequencing (ChIP-seq) of Pol V with an antibody that targets the largest and unique subunit of the Pol V complex (NRPE1) in wild type, *morchex* (*morc1 morc2 morc4 morc5 morc6 morc7*), and *mom1* mutants. To compare MORC/MOM1-mediated regulation with SUVH2/9- and non-CG DNA methylation–mediated Pol V occupancy, we also profiled Pol V in *suvh29* (*suvh2 suvh9*) and *ddcc* (*drm1 drm2 cmt2 cmt3*) mutants (Fig. 1a). Consistent with previous results revealing SUVH2/9 as the dominant determinants of Pol V binding^14^, nearly all wild type Pol V peaks were almost completely lost in the *suvh29* mutant (Fig. 1b). In contrast, in *mom1*, *morchex*, and *ddcc* mutants, Pol V peaks were often retained but also redistributed across the genome, with stronger redistributions detected in *ddcc*, followed by *mom1* and *morchex*. We identified differentially bound Pol V regions between wild type and mutant backgrounds, revealing 1,602, 1,677, 7,411, and 12,223 loci with reduced Pol V occupancy in *morchex*, *mom1*, *ddcc*, and *suvh29* mutants, respectively. Conversely, 2,785, 2,930, 10,313, and 22,264 regions exhibited significantly increased Pol V binding in these mutants (Fig. 1c). The substantial redistribution observed in *suvh29*, *morchex*, *mom1*, and *ddcc* mutants indicates that SUVH2/9, MORC, MOM1, and DNA methylation function to refine and spatially constrain Pol V chromatin distribution.

**Fig 1.**
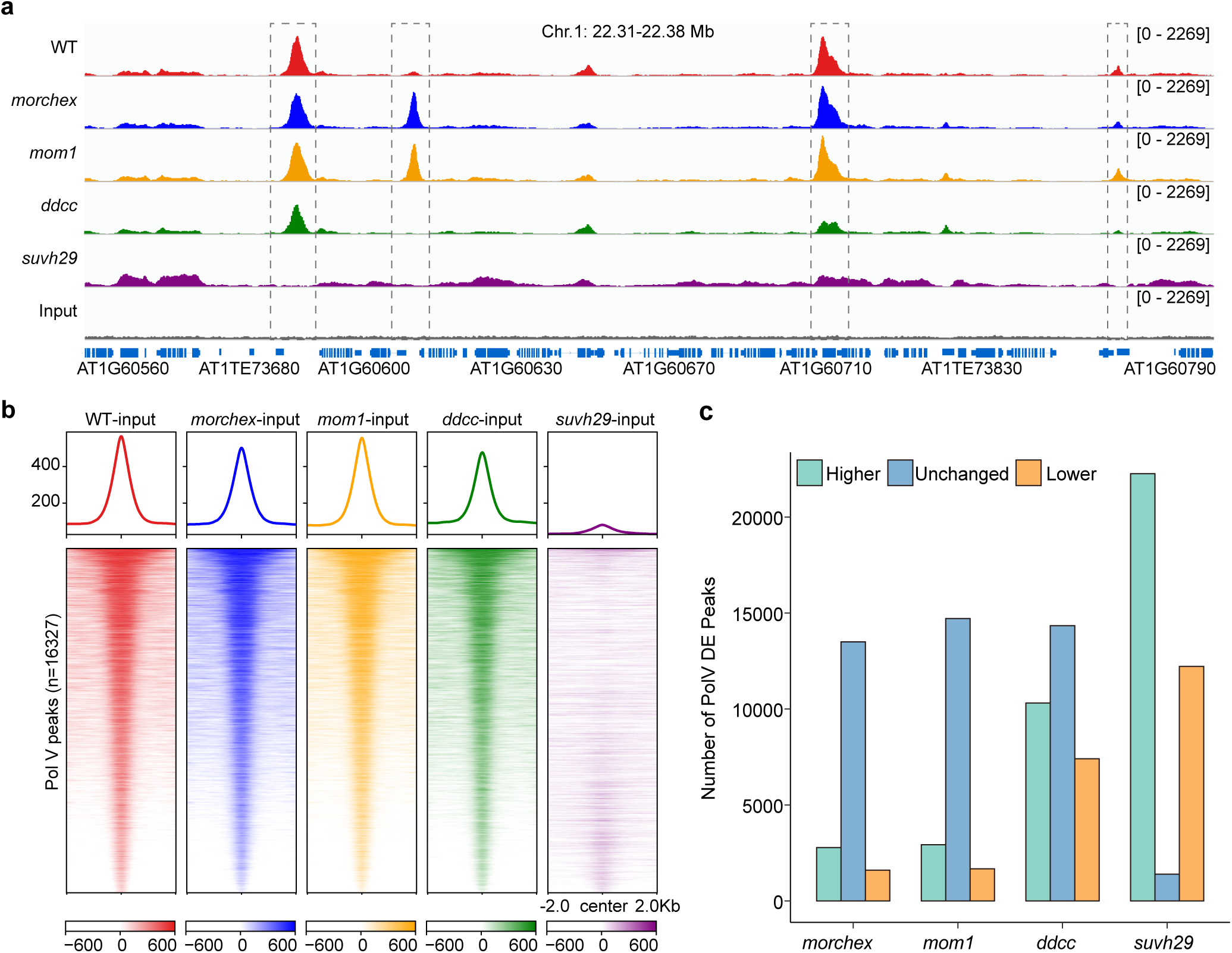
The essential role of chromatin regulators in regulating RNA polymerase V (Pol V) chromatin distribution. **a.** Representative screenshots showing the Pol V signals in wild type (WT), *morchex*, *mom1*, *ddcc*, and *suvh29* mutants. **b.** Metaplot showing input-normalized Pol V signal in WT and mutant backgrounds across WT-defined Pol V peaks (n = 16,327). **c.** Quantification of differentially bound Pol V regions in *morchex*, *mom1*, *ddcc*, and *suvh2/9* mutants relative to WT. Regions are classified as Higher (increased Pol V binding in mutants), Unchanged (no significant difference), or Lower (reduced Pol V binding in mutants).

### MORC and MOM1 synergistically affect Pol V occupancy

The MOM1 complex associates with RdDM target loci and facilitates MORC recruitment through a direct interaction between PIAL2 and MORC6 (ref^26^), suggesting that MOM1 and MORC function in a coordinated manner to regulate Pol V chromatin occupancy. To test this hypothesis, we generated higher-order mutants combining *morchex* and *mom1*, and profiled Pol V occupancy in this genetic background (Fig. 2a). UpSet analysis of differentially bound Pol V regions revealed both shared and distinct contributions of MORC and MOM1 to Pol V redistribution (Fig. 2b,c). For regions with increased Pol V occupancy, most peaks were unique to the *mom1 morchex* double mutant, with a substantially larger set size compared to either *morchex* or *mom1* single mutants. Only a limited fraction of hyper-bound regions overlapped between single mutants, indicating that MORC and MOM1 individually contribute to Pol V restriction but are insufficient alone to fully prevent its ectopic accumulation. In contrast, the *mom1 morchex* mutant exhibited a dramatic expansion of these regions, supporting a synergistic effect (Fig. 2b). We observed a similar but less pronounced pattern for regions with reduced Pol V occupancy. While *morchex* and *mom1* mutants displayed partially overlapping sets of “Pol V lower” loci, the *mom1 morchex* mutant showed an increased number of unique and combined affected regions (Fig. 2c). The relatively higher overlap among Pol V lower regions compared to Pol V higher regions suggests that MORC and MOM1 act more redundantly in maintaining Pol V at canonical sites, but cooperatively in preventing its redistribution.

**Fig 2.**
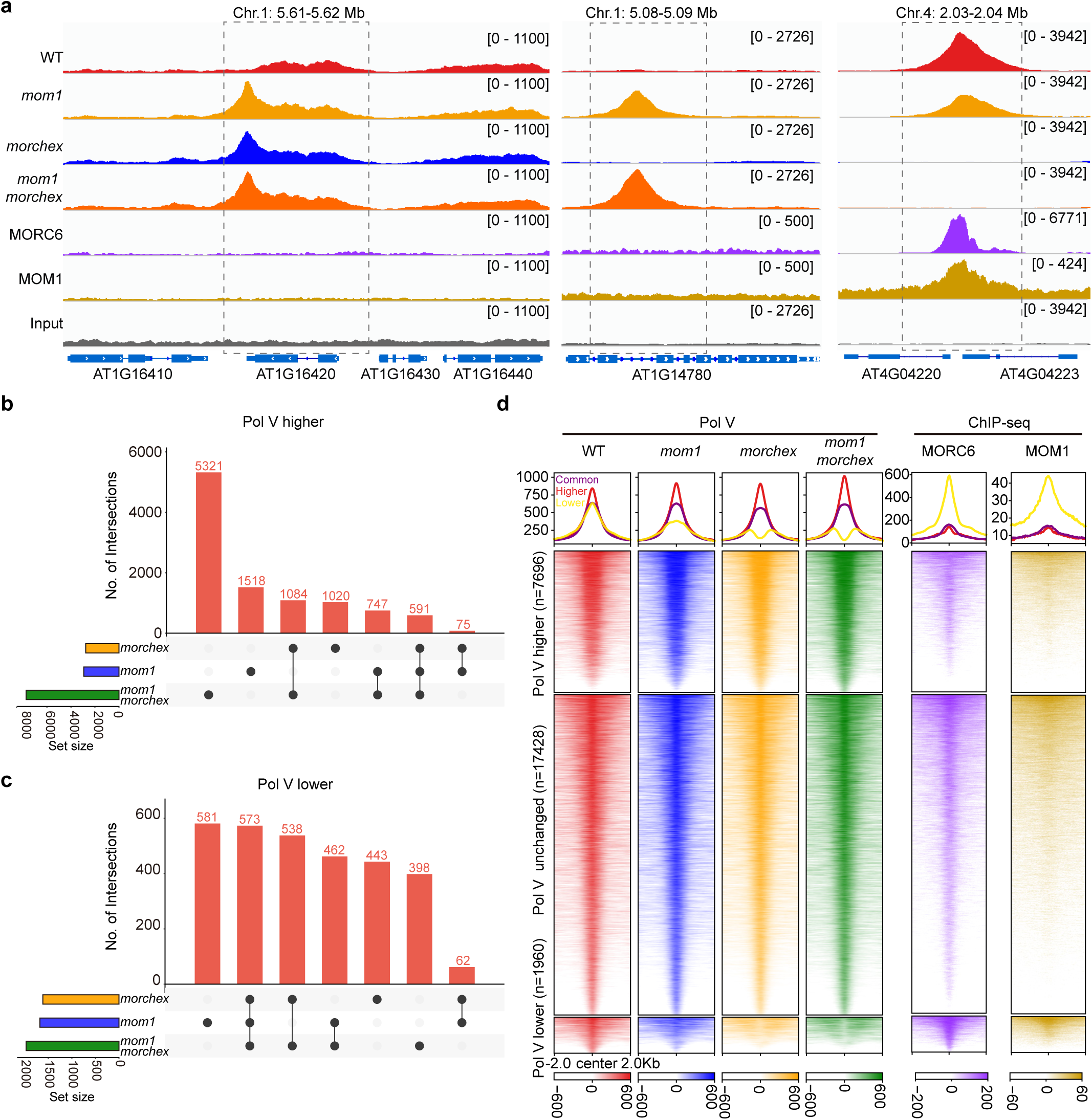
MORC and MOM1 synergistically affect Pol V chromatin distribution. **a.** Representative genome browser snapshots showing Pol V occupancy in wild type (WT), *morchex*, *mom1*, and *mom1 morchex* double mutants. The lower tracks display previously published ChIP-seq profiles of MORC6 and MOM1 at the same regions. **b.** Overlap of regions with higher Pol V binding in *morchex*, *mom1*, and *mom1 morchex* mutants. **c.** Overlap of regions with lower Pol V binding in *morchex*, *mom1*, and *mom1 morchex* mutants. **d.** Metaplot showing input-normalized Pol V signal together with MORC6 and MOM1 ChIP-seq signals across differentially bound Pol V regions identified in the *mom1 morchex* double mutant.

To further determine whether these redistributed Pol V sites are directly associated with MORC and MOM1, we examined previously published MORC6 and MOM1 ChIP-seq datasets^26^. Indeed, regions with lower Pol V signal in the *mom1 morchex* mutant were strongly enriched for both MORC6 and MOM1 occupancy, indicating that these loci represent bona fide targets of MORC and MOM1 (Fig. 2d). These results demonstrate that MORC and MOM1 are required to maintain proper Pol V occupancy at specific chromatin regions. The data also indicates that MORC contributes more prominently to regions with reduced Pol V binding, whereas MOM1 plays a broader role in constraining Pol V redistribution (Fig. 2d). This asymmetric contribution also further supports that MOM1 and MORC exert partially distinct but also overlapping functions in maintaining Pol V localization. Together, these findings support a model in which MORC and MOM1 act synergistically to spatially constrain Pol V, ensuring its proper distribution across the genome and preventing aberrant redistribution to ectopic sites.

### Chromatin and intrinsic sequence features underlie Pol V targeting

The redistribution of Pol V in the mutants demonstrates not only pivotal roles in determining Pol V chromatin binding but also provides a good opportunity to analyze previously uncharacterized Pol V transcriptional features. To this end, we investigated the genomic annotation of these Pol V binding loci in the *morchex* mutant. Pol V lower peaks exhibited a pronounced enrichment near promoter regions (Supplementary Fig. 1), whereas Pol V higher peaks and unchanged peaks displayed enrichment at distal intergenic regions (Supplementary Fig. 1). Furthermore, Pol V lower peaks were found in closer proximity to transcription start sites (TSSs) of genes exhibiting higher chromatin accessibility (Fig. 3a,b). Both the Pol V unchanged peaks and Pol V higher peaks were more enriched over TEs, with Pol V higher peaks showing a higher enrichment in long TEs (Fig. 3c). In line with a hypothesis of Pol V redistribution from active to inactive regions in the *morchex* mutant, the flanking regions of Pol V lower peaks exhibited lower TE content (Fig. 3c). The Pol V lower peaks exhibited a significantly higher enrichment of MORC signals, accompanied by higher levels of ROS1 and lower levels of H3.1 and 5mC methylation than the Pol V higher peaks (Fig. 3d-j). Thus, regions characterized by elevated Pol V signal were more likely to correspond to heterochromatic regions, whereas those with diminished Pol V signal tended to possess higher transcription potential.

**Fig. 3.**
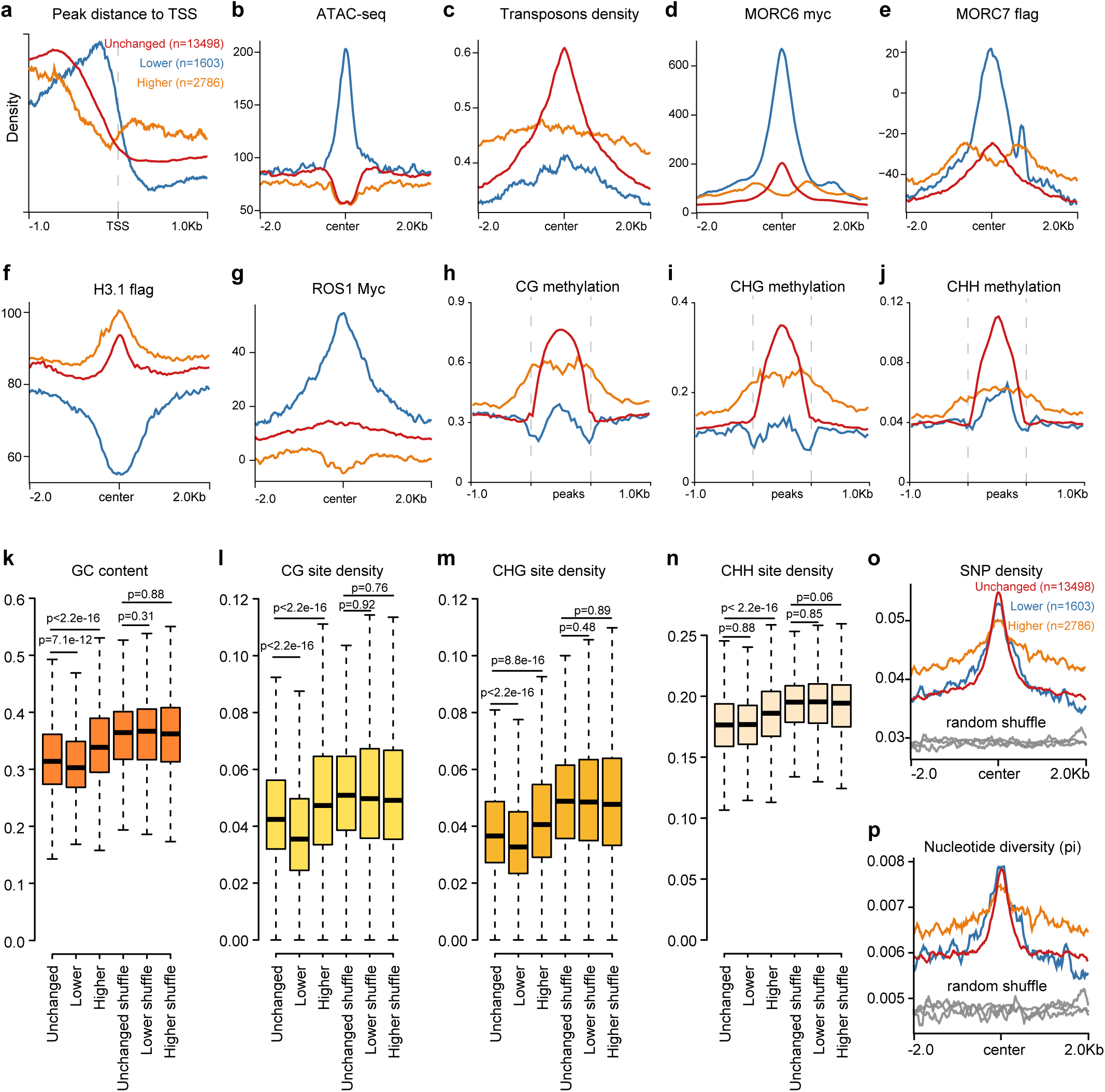
Chromatin and intrinsic sequence features associated with Pol V targeting. **a**. Density distribution of distances from Pol V peaks to transcription start sites (TSS). **b–g**. Metaplots showing chromatin features across Pol V–lower, Pol V–higher, and Pol V–unchanged peaks in the *morchex* mutant, including ATAC–seq signal (**b**), transposable element (TE) density (**c**), MORC6 ChIP–seq (**d**), MORC7 ChIP–seq (**e**), histone variant H3.1 ChIP–seq (**f**), and ROS1 ChIP–seq (**g**). **h–j**. Metaplots of DNA methylation levels in CG (**h**), CHG (**i**), and CHH (**j**) contexts across the three classes of Pol V peaks. **k–n**. Sequence composition of Pol V–associated regions, including GC content (**k**) and cytosine context densities (CG, **l**; CHG, **m**; CHH, **n**). **o,p**. SNP density (**o**) and nucleotide diversity (**p**) across Pol V–lower, Pol V–higher, and Pol V–unchanged peaks. For **k–n**, boxplots indicate median, interquartile range, and data distribution; statistical significance was assessed using two-sided Student’s *t*-test. Randomly shuffled genomic regions were used as controls (grey lines in metaplots).

We next investigated whether intrinsic DNA sequence features contribute to Pol V localization in the *morchex* mutant. Our analysis revealed a notable trend of lower overall GC content within Pol V binding regions across the genome (Fig. 3k). Pol V lower peaks exhibited the lowest GC content, while Pol V unchanged peaks displayed an intermediate level of GC content (Fig. 3k). Pol V higher peaks showed the highest GC content, indicating that MORC facilitates the recruitment of Pol V to regions with low GC content, or, in other words, AT-rich regions. Concomitantly, we assessed the densities of CG, CHG, and CHH (both methylated and unmethylated) sites within Pol V binding sites and observed relatively lower densities of these sites in Pol V binding regions, particularly in Pol V lower peaks (Fig. 3l-n). Furthermore, we calculated SNP density and nucleotide diversity of Pol V binding peaks by incorporating population genomic data of the *Arabidopsis* 1001 accessions^30^. Pol V exhibited a preference for binding to regions with higher genetic diversity in the *Arabidopsis* population (Fig. 3o,p). Together, these findings indicate that Pol V targeting is associated with a combination of chromatin context, DNA sequence composition, and genetic variability. Rather than being solely directed by DNA methylation, Pol V preferentially associates with AT-rich, sequence-diverse regions, while chromatin regulators such as MORC and MOM1 act to constrain its distribution within specific epigenetic domains.

### Proper Pol V distribution is reinforced by non-CG methylation

To dissect the relationship between DNA methylation and Pol V occupancy, we began by examining the methylation patterns at Pol V–associated loci in the published WGBS data of *suvh2/9* mutant^31^. At Pol V–unchanged regions in the *suvh2/9* mutant, CG, CHG, and CHH methylation levels remained relatively stable despite the loss of SUVH2/9, indicating that maintenance of DNA methylation at these loci is largely independent of SUVH2/9 (Fig. 4a). Regions exhibiting increased Pol V binding in *suvh2/9* showed a concomitant increase in DNA methylation across all sequence contexts, whereas regions with reduced Pol V occupancy displayed a corresponding decrease in DNA methylation (Fig. 4b,c). This coordinated change suggests that once Pol V is present at a locus, it is sufficient to initiate *de novo* DNA methylation, even in the absence of SUVH2/9 (Fig. 4d). These observations indicate that SUVH2/9 primarily functions to anchor Pol V at canonical RdDM loci, rather than being strictly required for methylation establishment per se.

**Fig. 4.**
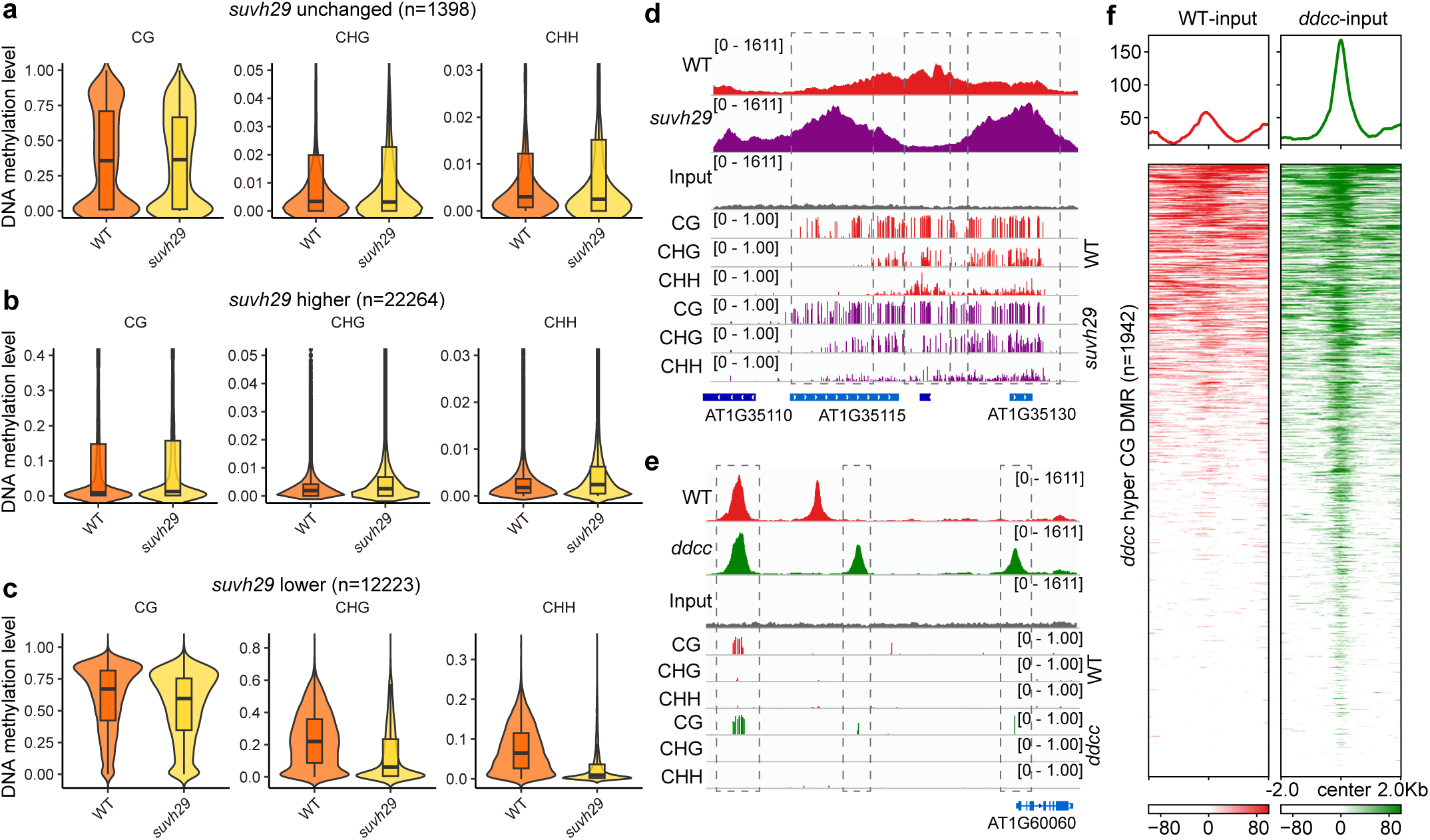
DNA methylation dynamics associated with Pol V redistribution and loss of SUVH2/9 function. **a–c**. Violin plots showing DNA methylation levels in CG, CHG, and CHH contexts at Pol V–associated loci in wild type (WT) and *suvh2/9* mutants. Loci are categorized based on Pol V changes in *suvh2/9* as unchanged (a, n = 1,398), higher (b, n = 12,223), and lower (c, n = 22,264). **d**. Representative genome browser view showing Pol V occupancy and DNA methylation profiles in WT and *suvh2/9*. Tracks include Pol V ChIP-seq signal, input control, and cytosine methylation levels (CG, CHG, CHH) in both genotypes. **e**. Representative genome browser view illustrating Pol V occupancy and DNA methylation profiles in WT and *ddcc* mutants. Despite loss of non-CG methylation, Pol V remains detectable at selected loci. **f**. Heatmaps and average profiles of input-normalized Pol V signal over CG hypermethylated differentially methylated regions (DMRs) in WT and *ddcc* (n = 1,942). Increased Pol V occupancy is observed at CG hyper-DMRs in *ddcc* relative to WT.

To further evaluate the role of non-CG methylation in Pol V targeting, we analyzed Pol V occupancy in the *ddcc* mutant, which lacks CHG and CHH methylation^32^. Despite the global loss of non-CG methylation, Pol V remained detectable at a subset of loci, particularly those enriched for CG methylation (Fig. 4e, Supplementary Fig. 2). Pol V occupancy was also markedly increased at CG hypermethylated regions in *ddcc* (Fig. 4e,f), suggesting that CG methylation alone can partially support Pol V association at specific genomic regions. Collectively, these results support a model in which DNA methylation and Pol V occupancy are not strictly interdependent. Instead, SUVH2/9-mediated recognition primarily determines the canonical positioning of Pol V, while Pol V itself can drive *de novo* methylation at newly occupied loci. Furthermore, non-CG methylation acts to reinforce and stabilize Pol V distribution, whereas CG methylation can partially compensate for its loss, enabling Pol V targeting in a methylation context-dependent manner.

### CMT3-mediated DNA methylation maintenance is influenced by Pol V redistribution

CMT3 plays a critical role in maintaining DNA methylation, particularly in the CHG context within regions that contain transposons and repetitive elements^32,34,35^. To investigate the impact of Pol V redistribution on DNA methylation maintenance, we conducted ChIP-seq experiments using pCMT3:CMT3-3xFLAG transgenic plants in both wild type and *morchex* mutant. In line with previous research, CMT3 displayed high enrichment within the body regions of long TEs (Supplementary Fig. 3)^35^. Consistent with the established role of CMT3 as a key CHG methylation maintenance enzyme functioning in coordination with H3K9me2, we detected substantial CMT3 enrichment at both Pol V unchanged and Pol V higher regions (Fig. 5a). In contrast, CMT3 occupancy was markedly lower at Pol V lower regions but remained detectable, suggesting these regions are mainly RNA directed DNA methylation sites. In the *morchex* mutant, CMT3 binding was higher at TEs, particularly over long TEs (Fig. 5b). For *morchex* Pol V lower sites, CMT3 binding was significantly reduced (p value < 2.16e-16, Fig. 5c-d), indicating that MORC activity contributes to sustaining CMT3 association at these loci. A likely explanation is that lower Pol V at these regions leads to reduced RdDM activity and associated CMT3-dependent maintenance methylation, as it is known that at some RdDM loci CMT3 and the RdDM machinery play overlapping roles in controlling both CHH and CHG methylation^36^.

**Fig. 5.**
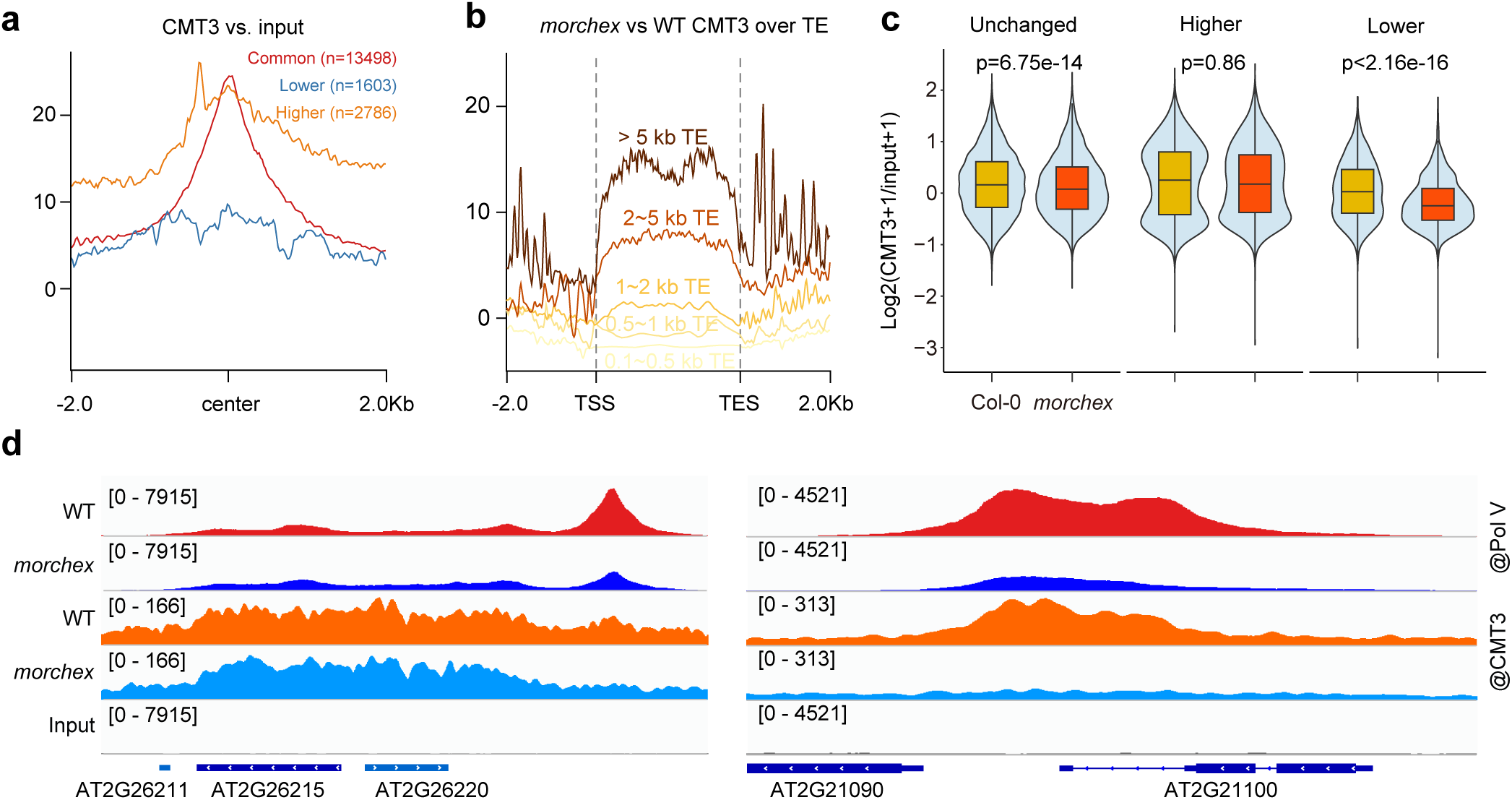
CMT3 binding is affected by redistribution of Pol V. **a.** Metaplot of wild-type CMT3 ChIP-seq over Pol V lower, Pol V higher, and Pol V unchanged peaks. **b.** Metaplot showing CMT3 ChIP-seq signal (log2 FLAG/input) over transposons grouped by lengths. TEs are divided into five groups: 0.1 to 0.5 kb TEs (n=12,018), 0.5 to 1 kb TEs (n=5,856), 1 to 2 kb TEs (n=3,471), 2 to 5 kb TEs (n=2,225), and > 5 kb TEs (n=1,022). **c.** A violin plot illustrates the binding levels of CMT3 over Pol V lower, Pol V higher, and Pol V unchanged peaks in wild-type and *morchex*. Boxplots within the violin plots display the upper quartile, median, and lower quartile of the dataset. P-values are computed using Student’s t-test. **d.** Representative genome browser snapshots showing Pol V and CMT3 occupancy in wild type (WT) and *morchex* mutants.

### Pol V transcription at ectopic sites triggers siRNA biogenesis and DNA methylation

To assess whether changes in Pol V binding led to changes in Pol V transcription, we conducted RNA immunoprecipitation sequencing (RIP-seq) in floral samples of wild-type and *morchex* mutant. Our analysis revealed distinct Pol V transcription across different genomic regions. Specifically, in Pol V unchanged peaks of the *morchex* mutant, we observed only subtle effects on Pol V transcription levels (Fig. 6a), suggesting a relatively modest alteration in its transcription activity. However, in Pol V higher regions, Pol V transcription levels significantly increased (p value = 3.611e-12, Fig. 6a,b). Conversely, in Pol V lower regions, Pol V transcription levels significantly decreased (p value = 3.1e-07, Fig. 6a,c). Pol II binding remained unaffected and, consistent with previous findings that Pol V transcripts lack poly A^6^, we did not observe obvious changes in mRNA level detected by strand-specific RNA-seq (Supplementary Fig. 4a,b), which excludes the possibility that these RNA transcripts captured by Pol V RIP-seq are transcribed by Pol II^33^. These findings collectively show that the loss of MORC proteins not only influences genomic occupancy but also profoundly impacts Pol V transcription.

**Fig. 6.**
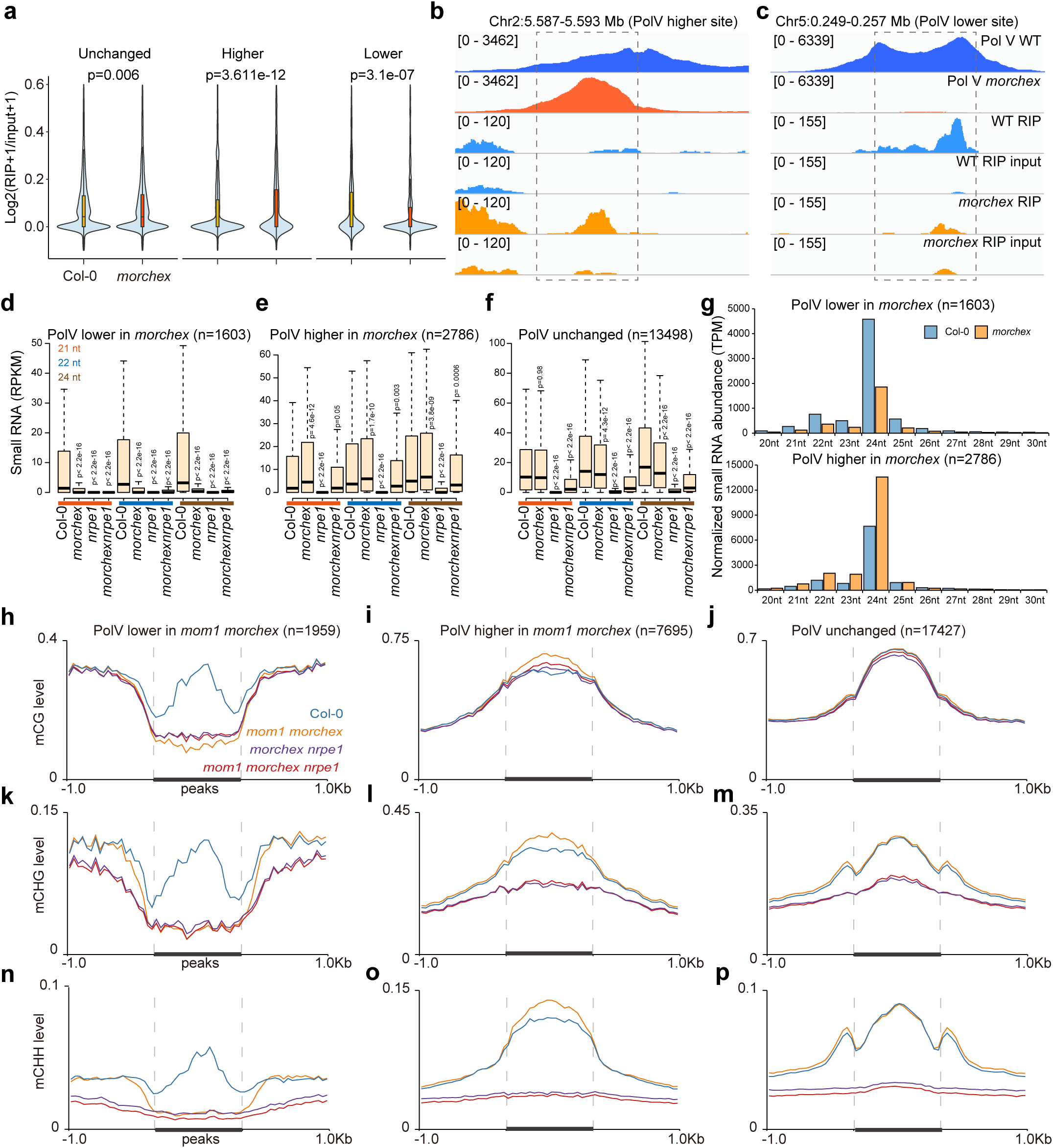
Redistributed Pol V initiates transcription, siRNA biogenesis, and DNA methylation. **a.** A violin plot illustrates the transcription levels of Pol V over Pol V lower, Pol V higher, and Pol V unchanged peaks. Boxplots within the violin plots display the upper quartile, median, and lower quartile of the dataset. P-values are computed using Student’s t-test. **b,c.** Representative screenshots from RIP-seq data depict the Pol V transcription patterns over Pol V lower and Pol V higher peaks. **d.** Boxplot displaying the abundance of 21-nt, 22-nt, and 24-nt small RNAs associated with Pol V lower peaks in Col-0, *morchex*, *nrpe1*, and *morchex nrpe1* mutants. **e.** Boxplot displaying the abundance of 21-nt, 22-nt, and 24-nt small RNAs associated with Pol V higher peaks in Col-0, *morchex*, *nrpe1*, and *morchex nrpe1* mutants. **f.** Boxplot displaying the abundance of 21-nt, 22-nt, and 24-nt small RNAs associated with Pol V unchanged peaks in Col-0, *morchex*, *nrpe1*, and *morchex nrpe1* mutants. Boxplots represent the upper quartile, median, and lower quartile of the dataset, with P-values calculated using Student’s t-test. **g.** small RNA length distribution of Pol V lower and higher peaks in Col-0 and *morchex* mutants. CG methylation level over Pol V lower (**h**), Pol V higher (**i**), and Pol V unchanged peaks (**j**) in Col-0, *mom1 morchex*, *morchex npre1*, and *mom1 morchex nrpe1* mutants. CHG methylation level over Pol V lower (**k**), Pol V higher (**l**), and Pol V unchanged (**m**) peaks in Col-0, *mom1 morchex*, *morchex npre1*, and *mom1 morchex nrpe1* mutants. CHH methylation level over Pol V lower (**n**), Pol V higher (**o**), and Pol V unchanged (**p**) peaks in Col-0, *mom1 morchex*, *morchex npre1*, and *mom1 morchex nrpe1* mutants.

As scaffold RNAs in guiding downstream AGO-siRNA complexes, Pol V transcripts often have a uracil (U) at position 10, which pairs complementarily with the adenine (A) at position 1 present in AGO4-bound 24-nt siRNAs^17^. To investigate whether the newly transcribed Pol V transcripts can initiate siRNA biogenesis, we conducted small RNA sequencing (small RNA-seq) in Col-0 (wild type) and *morchex* mutant. Levels of 21-nt, 22-nt, and 24-nt small RNAs were significantly (p value < 2.16e-16) reduced at Pol V lower sites, while their abundance increased at Pol V higher sites in the *morchex* mutant (Fig. 6 d-f). The 24-nt siRNAs were particularly affected by the redistribution of Pol V transcripts (Fig. 6g). Nucleotide bias analysis indicated that small RNA biogenesis at these sites initiated with an adenine (Supplementary Fig. 4c,d), consistent with the nucleotide bias typically observed in AGO loading small RNAs^37^. These findings suggest that the redistributed Pol V transcripts initiate siRNA biogenesis, with a significant impact on the abundance of small RNAs, particularly 24-nt siRNAs.

To determine whether the redistribution and altered transcriptional activity of Pol V leads to changes in DNA methylation, and whether these effects are Pol V dependent, we generated higher-order mutants by crossing *morchex* or *mom1 morchex* with *nrpe1*. We then performed whole-genome bisulfite sequencing (WGBS) in Col-0, *mom1 morchex*, *morchex nrpe1*, and *mom1 morchex nrpe1* mutants. Genome-wide changes in DNA methylation closely mirrored alterations in Pol V occupancy. Specifically, CG, CHG, and CHH methylation levels showed coordinated increases or decreases corresponding to gains or losses of Pol V occupancy in the *mom1 morchex* mutant (Fig. 6h–p). These methylation changes were largely abolished in the absence of NRPE1, indicating that they are dependent on Pol V activity. A similar pattern was observed in the *morchex* mutant across all categories of Pol V peaks (Supplementary Fig. 5). Together, these results demonstrate that redistribution of Pol V directly reshapes the DNA methylation landscape in an NRPE1-dependent manner, with MOM1 and MORC proteins playing an important role in mediating these effects.

## Discussion

RNA-directed DNA methylation (RdDM) has been viewed as a silencing pathway in plants, in which RNA Polymerase V (Pol V) is recruited to methylated regions through SUVH2/SUVH9-mediated recognition of pre-existing DNA methylation^31,38^. Here, we extend this model by demonstrating that Pol V chromatin positioning is dynamically regulated rather than passively dictated by DNA methylation or heterochromatic features. We show that the chromatin regulators MORC and MOM1 actively constrain Pol V to specific genomic regions, thereby shaping the specificity of RdDM. In the absence of this spatial constraint, Pol V is not simply lost but is redistributed into heavily methylated, CMT3-enriched heterochromatin, where it remains transcriptionally active, producing scaffold noncoding RNAs, promoting small RNA biogenesis, and reinforcing DNA methylation. These findings establish a combinatorial framework in which chromatin remodeling and epigenetic feedback collectively define Pol V distribution and function, positioning Pol V as an active determinant of epigenetic state rather than a passive downstream effector.

The results also support a hierarchical framework governing Pol V chromatin positioning. Consistent with previous studies, SUVH2 and SUVH9 act as primary determinants of Pol V recruitment by recognizing pre-existing DNA methylation, as their loss results in a near-complete depletion of Pol V occupancy from its normal locations^31^. However, this methylation-dependent recruitment alone is insufficient to fully define Pol V distribution. In the *ddcc* mutant, although non-CG methylation is largely abolished, Pol V remains detectable at a subset of loci, where CG methylation appears to partially support its association^32^. Regions exhibiting increased Pol V occupancy in *ddcc* also show elevated CG methylation, suggesting that Pol V can, in turn, promote CG methylation at these sites. Furthermore, redistributed Pol V remains functionally active even in the absence of SUVH2/9, MORC, or MOM1, as evidenced by its transcriptional activity and the establishment of DNA methylation at newly occupied loci. These observations indicate that the SUVH2/9–MORC–MOM1 regulatory system primarily governs the recruitment and spatial positioning of Pol V, rather than its catalytic activity. Instead, once localized, Pol V is sufficient to drive downstream epigenetic processes. These findings support a three-tier model in which methylation-dependent recruitment licenses Pol V binding, chromatin regulators refine its spatial distribution, and Pol V activity feeds back to shape DNA methylation patterns which in turn shape further Pol V recruitment.

Both MORC and MOM1 are evolutionarily conserved chromatin regulators implicated in chromatin compaction and transcriptional repression, and their loss leads to increased chromatin accessibility and altered transcription factor binding^19,21,24–27,39^. Previous studies have established roles for the DDR complex and SUVH2/9 in Pol V recruitment; however, our results place MORC and MOM1 as key regulators that act downstream to refine Pol V chromatin distribution^12,14,29^. MORC and MOM1 exhibit broad genomic localization across both repressive and accessible chromatin, suggesting that their function is not limited to heterochromatin but extends to actively shaping chromatin environments that lack canonical silencing marks^25,26,29^. This dual localization provides a mechanistic basis for their role in constraining Pol V to AT-rich, accessible regions, where targeting signals are otherwise weak and require additional regulatory input^27,31^. In this context, the redistribution of Pol V in *morchex* or *mom1* mutants from accessible chromatin to CMT3-enriched heterochromatin highlights a failure of spatial restriction rather than a change in intrinsic targeting preference.

Pol V preferentially binds to AT-rich regions, consistent with observations for Pol II and our recent structural evidence that A-rich DNA promotes Pol V transcription termination, suggesting that intrinsic sequence composition contributes to Pol V transcription dynamics and the establishment of DNA methylation landscapes^40,41^. Pol V occupancy is enriched at regions with high nucleotide diversity, raising the possibility that Pol V targeting is coupled to the detection of recently diverged or newly inserted TEs. Such coupling would enable Pol V to function as part of a surveillance system that monitors genome variation and facilitates the establishment of epigenetic silencing at emerging loci. Alternatively, it may simply be that these Pol V targeted regions are under less evolutionary pressure and thus diversify more over time.

Together, our results support a model in which MORC proteins and MOM1 play important roles in orchestrating precise Pol V chromatin occupancy (Fig. 7). In this model, Pol V exhibits a latent capacity to associate with methylation-rich heterochromatic regions, but its distribution is constrained by MORC–MOM1 to active chromatin environments, thereby ensuring proper targeting of the RdDM machinery. Loss of this spatial constraint leads to widespread redistribution of Pol V into heterochromatin, where it remains transcriptionally competent and reinforces DNA methylation and small RNA production. This regulatory mechanism is particularly important for RdDM at loci embedded within more active chromatin contexts, where targeting signals are inherently weaker. More broadly, our findings highlight spatial regulation of Pol V as a key determinant of DNA methylation landscapes, linking chromatin organization to epigenetic plasticity, genome stability, and adaptive responses.

**Fig. 7.**
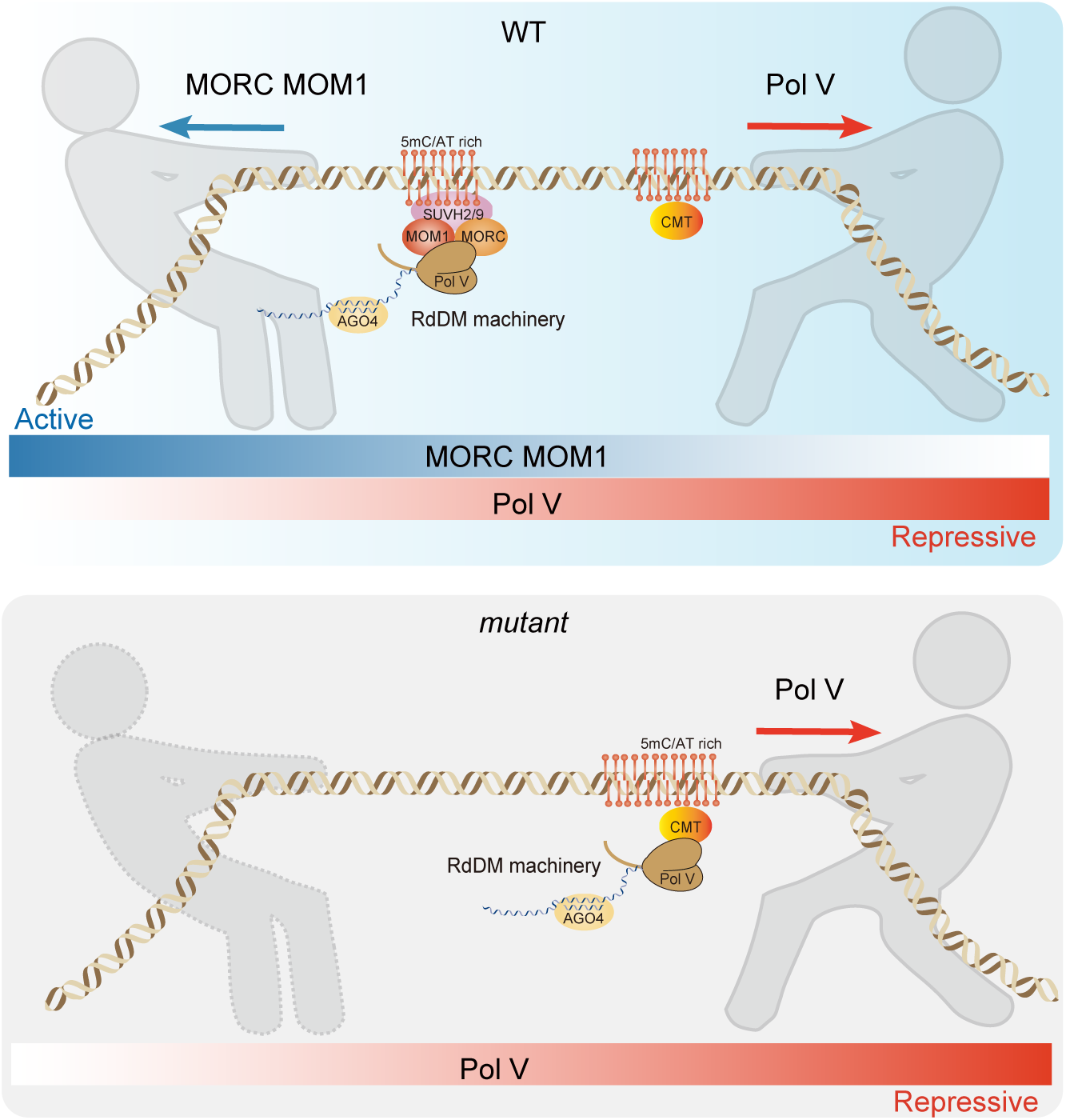
The proposed model of this study. The intrinsic characteristic of Pol V is a high affinity for enrichment within relatively silent heterochromatin regions. The presence of MORC and MOM1 proteins serves as a tether, empowering Pol V to occupy regions characterized by higher chromatin accessibility. This balance is essential for the proper establishment and propagation of RdDM sites, particularly within active genomic regions.

## Methods

### Plant materials and growth conditions

All plants included in this study were grown under standard greenhouse conditions, maintaining a temperature of 22 to 25 °C with a photoperiod of 16 hours of light and 8 hours of darkness. The following previously generated plant lines were used in this study: *ddcc* was generated by crossing *drm1* (SALK_031705), *drm2* (SALK_150863), *cmt2-7* (WISCDSLOX7E02) and *cmt2-3* (SALK_012874)^32^, *suvh2 suvh9* is was generated by crossing *suvh2* (Salk _079574.17.40) and *suvh9* (Salk_048033)^31^, *nrpe1-11* (SALK_029919)^42^, *morchex* was generated by crossing *morc1-2* (SAIL_893_B06), *morc2-1* (SALK_072774C), *morc4-1* (SALK_051729)*, morc5-1* (SALK_049050C), *morc6-3* (GABI_599B06), and *morc7-1* (SALK_051729)^43^.

### Epitope-tagged transgenic lines

Full-length genomic DNA fragments along with native promoter sequences of CMT3 were cloned into pENTR/D vectors (Invitrogen), and then transferred into modified destination vectors carrying 3xFLAG with LR Clonase (Invitrogen). For detailed information on the primers used in this study, please refer to **Supplementary Table 1**.

### ChIP-seq library preparation

2-4 g of flower tissue were harvested from 4- to 5-week-old plants, and ground with liquid nitrogen as described previously^12^. To fix the chromatin, a nuclei isolation buffer containing 1% formaldehyde was applied to resuspend tissue powders for 10 min at room temperature, and the cross-linking reaction was terminated using freshly prepared 2 M glycine. Shearing was performed with Bioruptor Plus (Diagenode) for 30 cycles (30 seconds on and 30 seconds off), and immunoprecipitations with antibodies were performed overnight at 4°C. The anti-FLAG M2 (Sigma) and Pol V antibody were used in this study. Magnetic Protein A and Protein G Dynabeads (Invitrogen) were added and incubated at 4°C for 2 h. The reverse crosslink was carried out overnight at 65°C. The protein-DNA mix was then digested with Protease K (Invitrogen) at 45°C for 4 h. The DNA was purified and precipitated with 3 M Sodium Acetate (Invitrogen), glycoBlue (Invitrogen) and Ethanol overnight at -20°C. The precipitated DNA was then used for library preparation using the Ovation Ultra Low System V2 kit (NuGEN), and then sequenced using an Illumina NovaSeq 6000 sequencer.

### RIP-seq library preparation

2-4 g of flower tissue were collected from 4- to 5-week-old plants and ground with liquid nitrogen^28^. The Honda buffer, containing 1% formaldehyde, was used to resuspend and fix the chromatin for 10 min at room temperature. Freshly prepared 2 M glycine was then used to terminate the cross-linking reaction, and filtered through two layers of Miracloth into a 50-ml Falcon and centrifuged at 3000*g* for 8 min at 4°C. The pellet containing nuclei was resuspended in 1 mL Honda buffer and centrifuged at 2500*g* for 5 min at 4°C. This washing step was repeated two times. The pellet was then resuspended in 500 μL Nuclei Lysis Buffer and sonicated on Bioruptor Plus (Diagenode) for 30 seconds on and 30 seconds off). The extract was then centrifuged for 10 min at 16,000*g* at 4°C, and the supernatant was transferred to a fresh tube. 100 μL of nuclei extract prepared from formaldehyde cross-linked seedlings was diluted with 900 μL of ChIP dilution buffer (1.1% Triton X-100, 1.2 mM EDTA, 16.7 mM Tris-HCl, pH 8, 167 mM NaCl, and 160 units/mL RnaseOUT) and centrifuged at 16,000*g* for 10 min at 4°C. The supernatant was transferred to Eppendorf tube, and 10 μL of the supernatant was saved as RNA inputs. 40 μL protein A dynabeads for each RIP sample was washed 5 times with 1 mL of binding/washing buffer. The beads were resuspended in 50 μL of binding/washing buffer. The beads and 1 μg of Pol V antibody were added to the diluted nuclei extract, and the samples were incubated on a rotator for 3 h at 4°C. The beads were then washed with 1 mL binding/washing buffer containing 40 units/ml RnaseOUT six times. For each wash, the beads were kept on a rotator at 4°C for 5 min and centrifuged at 2000*g* for 2 min, and the supernatant was then removed. To elute the protein-RNA complexes, 60 μL of RIP elution buffer (100 mM Tris HCl, pH 8.0, 10 mM EDTA, 1% SDS, and 800 units/mL RnaseOUT) was added to the beads, and the tubes were incubated on a rotator at room temperature for 10 min. The beads were centrifuged at 4000*g* for 1 min, and the supernatant was saved. The elution was repeated with an additional 60 μL of RIP elution buffer at 65°C with mixing (800 rpm) for 10 min. The samples were centrifuged and the supernatants from two elution were combined. 110 μL of RIP elution buffer was added to the 10 μL of RNA input sample. 1.2 μL of 20 mg/mL Proteinase K was added to each RIP or input sample and incubated at 65°C for 1 h. The RNA was isolated using Trizol reagent (Zymo) following the manufacturer’s instructions. Finally, library preparation was performed for mutant and wild-type samples by TruSeq Stranded mRNA kit (Illumina). mRNA or rRNA depletion steps were skipped and the heat fragmentation replaced with an incubation on ice for 5 min.

### Whole-genome bisulfite sequencing (WGBS) library preparation

Flower tissues were used for WGBS libraries preparation. Genomic DNA was extracted and converted with bisulfite treatment with EpiTect Bisulfite Kit (Qiagen), following the manufacturer’s instructions. Sonicated genomic DNA from samples were end-repaired and ligated with TruSeq DNA single adapters (Illumina) using a Kapa DNA HyperPrep kit (Roche) and converted with an EpiTect Bisulfite Kit (Qiagen). Converted DNA fragments were amplified by MyTaq polymerase (Bioline) for 12 cycles. The libraries were run on D1000 ScreenTape (Agilent) to determine the quality and size and purified by AMPure XP beads (Beckman Coulter). The purified DNA was used for library preparation using the Ovation Ultra Low System V2 kit (NuGEN), and then sequenced using an Illumina NovaSeq 6000 sequencer.

### Small RNA-seq analysis

Total RNA from floral tissues of wildtype and mutant plants extracted with the Direct-zol RNA Miniprep kit (Zymo) were used for small RNA sequencing. Two micrograms of total RNA mixed with the 2x RNA loading dye (NEB) were denatured at 65°C for 10 min and immediately chilled on ice. Then the denatured total RNA was separated on 15% TBE urea gel (Invitrogen) and small RNAs with size from 15 to 40 nt were purified for small RNA libraries. NEBNext Small RNA Library Prep Set for Illumina (Multiplex Compatible) (NEB) were used for small RNA library generation according to the manufacturer’s instructions. Adaptor sequence (TGGAATTCTCGG) was trimmed with trim_galore, and trimmed reads were mapped to the reference genome TAIR10 using Bowtie2 with only one unique hit and zero mismatches^44^.

### RNA-seq analysis

Cleaned short reads were aligned to the reference genome, TAIR10, by Bowtie2 (v2.1.0)^44^. Expression abundance was then calculated by RSEM using the default parameters^45^. Heatmaps were visualized using the R package pheatmap^46^. Differential expression analysis was conducted using edgeR^47^. A threshold of p-value < 0.05 and Fold Change > 2 were used to decide whether there were any significant differences in expression between samples.

### ChIP-seq analysis

ChIP-seq data was aligned to the TAIR10 reference genome with Bowtie2 (v2.1.0)^44^, only including uniquely mapped reads without any mismatches. Duplicated reads were removed by Samtools (v1.9)^48^. ChIP-seq peaks were called by MACS2 (v2.1.1) and annotated using ChIPseeker^49^. Differential peaks were called by the bdgdiff function in MACS2^50^. Differential peaks identified in this study is available at Supplementary Table 2. ChIP-seq data metaplots were plotted by deeptools (v2.5.1)^51^. Previously published MORC6, MORC7, and H3.1 ChIP-seq data were included in this analysis^25,26,52^.

### RIP-seq analysis

Cleaned short reads were aligned to the reference genome TAIR10 by Bowtie2 (v2.1.0) with default settings^44^. Duplicated reads were removed by Samtools. The bamcoverage function in deeptools (v2.5.1) was used to convert mapping files into track files and visualize in Integrative Genomics Viewer (IGV, v2.9.2)^53,54^.

### WGBS analysis

Trim_galore (http://www.bioinformatics.babraham.ac.uk/projects/trim_galore/) was used to trim adapters in the WGBS reads. Then the trimmed reads were aligned to the TAIR10 reference genome by BSMAP (v2.90), allowing two mismatches and one best hit (-v 2 -w 1)^55^. Reads with three or more consecutive CHH sites were considered to be unconverted reads and were filtered out. DNA methylation levels of each cytosine were defined as mC/ (#C + #T).

## Supplementary Information

The online version contains supplementary material available at xxxx.

## Authors’ contributions

SEJ and ZZ conceived the study. XH, ZL, and YX performed experiments. SF performed high throughput sequencing. XH and ZZ analyzed the data. XH, ZZ, and SEJ wrote the manuscript with help from all authors. All authors approved the final version of the manuscript and agreed on the content and conclusions.

## Acknowledgements

We thank M. Akhavan and the UCLA BSCRC High Throughput BioSequencing Core for their technical assistance. We thank Life Science Editors for editing services. The work in the Zhong laboratory was financially supported by the Natural Science Foundation of Sichuan Province (2026NSFSCZY0075, 26QNJJB0389), the Fundamental Research Funds for the Central Universities. The work in the Jacobsen laboratory was supported by a grant from the W.M. Keck Foundation. S.E.J. is an investigator of the Howard Hughes Medical Institute.

## Availability of data and materials

Data supporting the findings of this work are available within the paper and its Supplementary Information files. All high-throughput sequencing data generated in this study are accessible at NCBI’s Gene Expression Omnibus (GEO) via GEO Series accession number GSE342837 (https://www.ncbi.nlm.nih.gov/geo/query/acc.cgi?acc=GSE342837).

## Supplementary Figures

**Supplementary Fig. 1** Genomic distribution enrichment of Pol V lower (n=1603) (**a**), Pol V higher (n=2786) (**b**), and Pol V unchanged peaks (n=13498) (**c**).

**Supplementary Fig. 2** Violin plots showing DNA methylation levels in CG, CHG, and CHH contexts at Pol V–associated loci in wild type (WT) and *ddcc* mutants. Loci are categorized based on Pol V changes in *ddcc* as unchanged (**a**, n = 14,342), higher (**b**, n = 10,313), and lower (**c**, n = 7,411).

**Supplementary Fig. 3 CMT3 binding over TEs.** Metaplot showing CMT3 ChIP-seq signal (log2 FLAG/input) over transposons grouped by lengths. TEs are divided into five groups: 0.1 to 0.5 kb TEs (n=12,018), 0.5 to 1 kb TEs (n=5,856), 1 to 2 kb TEs (n=3,471), 2 to 5 kb TEs (n=2,225), and > 5 kb TEs (n=1,022).

**Supplementary Fig. 4 a.** Violin plot illustrating Pol II binding level over Pol V lower, Pol V higher, and Pol V unchanged peaks. **b.** Violin plot displaying poly-A RNA transcription levels over Pol V lower, Pol V higher, and Pol V unchanged peaks. **f-g.** Nucleotide bias of small RNA of Pol V lower, Pol V higher, and Pol V unchanged peaks in Col-0 and *morchex* mutants.

**Supplementary Fig. 5 DNA methylation dynamics associated with Pol V redistribution in *morchex* mutant.** CG methylation level over Pol V lower (**a**), Pol V higher (**b**), and Pol V unchanged peaks (**c**) in WT, *morchex*, *npre1*, and *nrpe1 morchex* mutants. CHG methylation level over Pol V lower (**d**), Pol V higher (**e**), and Pol V unchanged (**f**) peaks in WT, *morchex*, *npre1*, and *nrpe1 morchex* mutants. CHH methylation level over Pol V lower (**g**), Pol V higher (**h**), and Pol V unchanged (**i**) peaks in WT, *morchex*, *npre1*, and *nrpe1 morchex* mutants.

## Supplementary Tables

**Supplementary Table 1** Primers used in this study.

**Supplementary Table 2** List of Pol V differential peaks between Col-0 and mutants.

## References

1. Kwak, H., Fuda, N. J., Core, L. J. & Lis, J. T. Precise maps of RNA polymerase reveal how promoters direct initiation and pausing. Science (1979). 339, 950–953 (2013).

2. Vermulst, M. et al. Transcription errors induce proteotoxic stress and shorten cellular lifespan. Nat. Commun. 6, 8065 (2015).

3. Matsui, T., Segall, J., Weil, P. A. & Roeder, R. G. Multiple factors required for accurate initiation of transcription by purified RNA polymerase II. Journal of Biological Chemistry 255, 11992–11996 (1980).

4. Xie, G. et al. Structure and mechanism of the plant RNA polymerase V. Science (1979). 379, 1209–1213 (2023).

5. Tucker, S. L., Reece, J., Ream, T. S. & Pikaard, C. S. Evolutionary History of Plant Multisubunit RNA Polymerases IV and V Subunit Origins via Genome-Wide and Segmental Gene Duplications, Retrotransposition, and Lineage-Specific Subfunctionalization. in Cold Spring Harbor symposia on quantitative biology sqb-2010 (Cold Spring Harbor Laboratory Press, 2011).

6. Wierzbicki, A. T., Haag, J. R. & Pikaard, C. S. Noncoding transcription by RNA polymerase Pol IVb/Pol V mediates transcriptional silencing of overlapping and adjacent genes. Cell 135, 635–648 (2008).

7. Wierzbicki, A. T., Ream, T. S., Haag, J. R. & Pikaard, C. S. RNA polymerase V transcription guides ARGONAUTE4 to chromatin. Nat Genet 41, 630 (2009).

8. Haag, J. R. & Pikaard, C. S. Multisubunit RNA polymerases IV and V: purveyors of non-coding RNA for plant gene silencing. Nat. Rev. Mol. Cell Biol. 12, 483–492 (2011).

9. Zhong, X. et al. Molecular mechanism of action of plant DRM de novo DNA methyltransferases. Cell 157, 1050–1060 (2014).

10. Marasco, M., Li, W., Lynch, M. & Pikaard, C. S. Catalytic properties of RNA polymerases IV and V: accuracy, nucleotide incorporation and rNTP/dNTP discrimination. Nucleic Acids Res. 45, 11315–11326 (2017).

11. Matzke, M. A., Kanno, T. & Matzke, A. J. M. RNA-directed DNA methylation: the evolution of a complex epigenetic pathway in flowering plants. Annu. Rev. Plant Biol. 66, 243–267 (2015).

12. Zhong, X. et al. DDR complex facilitates global association of RNA polymerase V to promoters and evolutionarily young transposons. Nat. Struct. Mol. Biol. 19, 870–875 (2012).

13. Tsuzuki, M. et al. Broad noncoding transcription suggests genome surveillance by RNA polymerase V. Proceedings of the National Academy of Sciences 117, 30799–30804 (2020).

14. Liu, Z.-W. et al. The SET domain proteins SUVH2 and SUVH9 are required for Pol V occupancy at RNA-directed DNA methylation loci. PLoS Genet. 10, e1003948 (2014).

15. Zhong, X. et al. Domains rearranged methyltransferase3 controls DNA methylation and regulates RNA polymerase V transcript abundance in Arabidopsis. Proceedings of the National Academy of Sciences 112, 911–916 (2015).

16. Law, J. A. et al. A protein complex required for polymerase V transcripts and RNA-directed DNA methylation in Arabidopsis. Current Biology 20, 951–956 (2010).

17. Liu, W. et al. RNA-directed DNA methylation involves co-transcriptional small-RNA-guided slicing of polymerase V transcripts in Arabidopsis. Nat Plants 4, 181 (2018).

18. Harris, C. J., Zhong, Z., Ichino, L., Feng, S. & Jacobsen, S. E. H1 restricts euchromatin-associated methylation pathways from heterochromatic encroachment. Elife 12, RP89353 (2024).

19. Dong, W. et al. MORC domain definition and evolutionary analysis of the MORC gene family in green plants. Genome Biol. Evol. 10, 1730–1744 (2018).

20. Kim, H. et al. The Gene-Silencing Protein MORC-1 Topologically Entraps DNA and Forms Multimeric Assemblies to Cause DNA Compaction. Mol. Cell 75, 700–710.e6 (2019).

21. Pastor, W. A. et al. MORC1 represses transposable elements in the mouse male germline. Nat. Commun. 5, 5795 (2014).

22. Li, S. et al. Mouse MORC3 is a GHKL ATPase that localizes to H3K4me3 marked chromatin. Proc. Natl. Acad. Sci. U. S. A. 113, E5108–E5116 (2016).

23. Desai, V. P. et al. The role of MORC3 in silencing transposable elements in mouse embryonic stem cells. Epigenetics Chromatin 14, 1–14 (2021).

24. Moissiard, G. et al. MORC family ATPases required for heterochromatin condensation and gene silencing. Science (1979). 336, 1448–1451 (2012).

25. Xue, Y. et al. Arabidopsis MORC proteins function in the efficient establishment of RNA directed DNA methylation. Nat. Commun. 12, 4292 (2021).

26. Li, Z. et al. The MOM1 complex recruits the RdDM machinery via MORC6 to establish de novo DNA methylation. Nat. Commun. 14, 4135 (2023).

27. Zhong, Z. et al. MORC proteins regulate transcription factor binding by mediating chromatin compaction in active chromatin regions. Genome Biol. 24, 96 (2023).

28. Böhmdorfer, G. et al. Long non-coding RNA produced by RNA polymerase V determines boundaries of heterochromatin. Elife 5, e19092 (2016).

29. Gallego-Bartolome, J. et al. Co-targeting RNA polymerases IV and V promotes efficient de novo DNA methylation in Arabidopsis. Cell 176, 1068–1082. e19 (2019).

30. Alonso-Blanco, C. et al. 1,135 Genomes Reveal the Global Pattern of Polymorphism in Arabidopsis thaliana. Cell 166, 481–491 (2016).

31. Sigman, M. J. et al. An siRNA-guided ARGONAUTE protein directs RNA polymerase V to initiate DNA methylation. Nat. Plants 7, 1461–1474 (2021).

32. Cao, X. et al. Role of the DRM and CMT3 methyltransferases in RNA-directed DNA methylation. Current biology 13, 2212–2217 (2003).

33. Stroud, H. et al. Non-CG methylation patterns shape the epigenetic landscape in Arabidopsis. Nat Struct Mol Biol 21, 64 (2014).

34. Du, J. et al. Dual binding of chromomethylase domains to H3K9me2-containing nucleosomes directs DNA methylation in plants. Cell 151, 167–180 (2012).

35. Wang, F., Johnson, N. R., Coruh, C. & Axtell, M. J. Genome-wide analysis of single non-templated nucleotides in plant endogenous siRNAs and miRNAs. Nucleic Acids Res. 44, 7395–7405 (2016).

36. Liu, Z. W. et al. Two Components of the RNA-Directed DNA Methylation Pathway Associate with MORC6 and Silence Loci Targeted by MORC6 in Arabidopsis. PLoS Genet. 12, e1006026 (2016).

37. Johnson, L. M. et al. SRA-and SET-domain-containing proteins link RNA polymerase V occupancy to DNA methylation. Nature 507, 124–128 (2014).

38. Kirshner, J. A. et al. Regulation of MORC-1 is key to the CSR-1–mediated germline gene licensing mechanism in C. elegans. Science Advances 11, (2025).

39. Geisberg, J. V et al. Nucleotide-level linkage of transcriptional elongation and polyadenylation. Elife 11, e83153 (2022).

40. Xie, G. et al. A spontaneous termination mechanism of RNA polymerase V shapes the DNA methylation landscape in plants. The EMBO Journal 2026 1–14 (2026) doi:10.1038/S44318-026-00763-7.

41. Pontier, D. et al. Reinforcement of silencing at transposons and highly repeated sequences requires the concerted action of two distinct RNA polymerases IV in Arabidopsis. Genes Dev 19, 2030–2040 (2005).

42. Harris, C. J., et al. Arabidopsis AtMORC4 and AtMORC7 Form Nuclear Bodies and Repress a Large Number of Protein-Coding Genes. PLoS Genet. 12, e1005998–e1005998 (2016).

43. Langmead, B. & Salzberg, S. L. Fast gapped-read alignment with Bowtie 2. Nat. Methods 9, 357–359 (2012).

44. Li, B. & Dewey, C. N. RSEM: accurate transcript quantification from RNA-Seq data with or without a reference genome. BMC Bioinformatics 12, 1–16 (2011).

45. Kolde, R. Pheatmap: pretty heatmaps. R package version 61, 617 (2012).

46. Robinson, M. D., McCarthy, D. J. & Smyth, G. K. edgeR: a Bioconductor package for differential expression analysis of digital gene expression data. bioinformatics 26, 139–140 (2010).

47. Li, H. et al. The Sequence Alignment/Map format and SAMtools. Bioinformatics 25, 2078–2079 (2009).

48. Yu, G., Wang, L.-G. & He, Q.-Y. ChIPseeker: an R/Bioconductor package for ChIP peak annotation, comparison and visualization. Bioinformatics 31, 2382–2383 (2015).

49. Zhang, Y. et al. Model-based Analysis of ChIP-Seq (MACS). Genome Biol. 9, R137 (2008).

50. Ramírez, F., Dündar, F., Diehl, S., Grüning, B. A. & Manke, T. deepTools: a flexible platform for exploring deep-sequencing data. Nucleic Acids Res. 42, W187–W191 (2014).

51. Zhong, Z. et al. Histone chaperone ASF1 mediates H3. 3-H4 deposition in Arabidopsis. Nat. Commun. 13, 6970 (2022).

52. Quinlan, A. R. & Hall, I. M. BEDTools: a flexible suite of utilities for comparing genomic features. Bioinformatics 26, 841–842 (2010).

53. Robinson, J. T. et al. Integrative genomics viewer. Nat. Biotechnol. 29, 24–26 (2011).

54. Xi, Y. & Li, W. BSMAP: whole genome bisulfite sequence MAPping program. BMC Bioinformatics 10, 232 (2009).

