## supplementary figure for "MORC and MOM1 spatially constrain RNA Polymerase V chromatin positioning to shape DNA methylation landscapes"

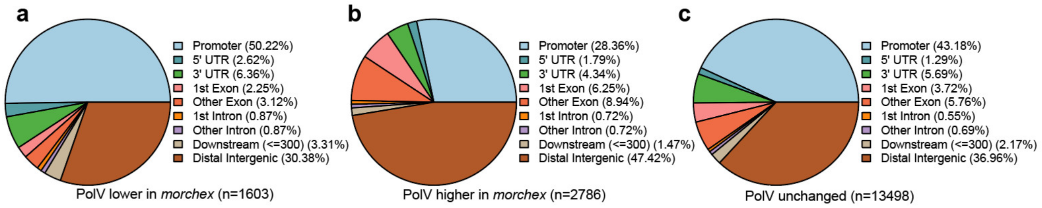


**Supplementary Fig. 1** Genomic distribution enrichment of Pol V lower (n=1603) (**a**), Pol V higher (n=2786) (**b**), and Pol V unchanged peaks (n=13498) (**c**).

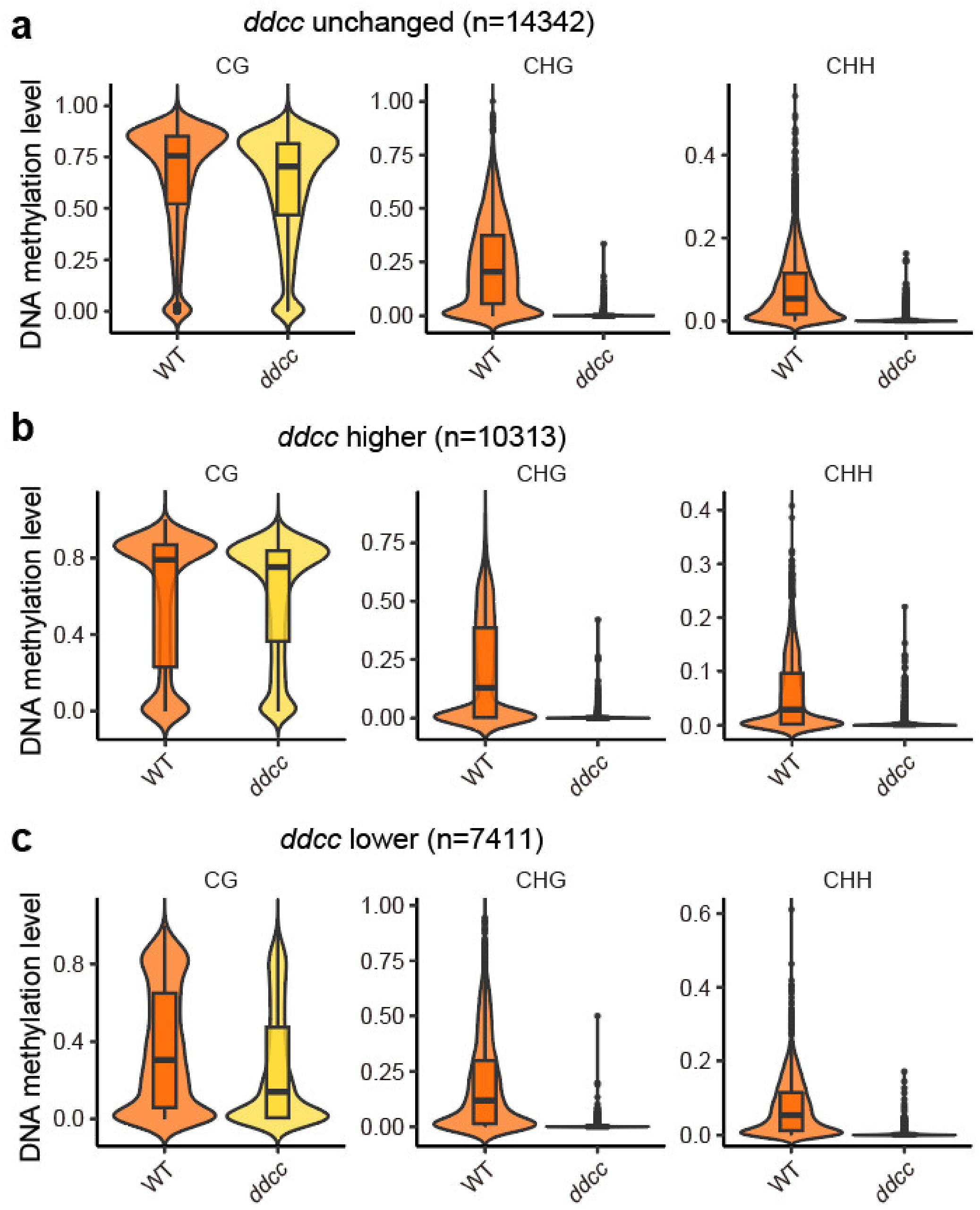


**Supplementary Fig. 2** Violin plots showing DNA methylation levels in CG, CHG, and CHH contexts at Pol V–associated loci in wild type (WT) and *ddcc* mutants. Loci are categorized based on Pol V changes in *ddcc* as unchanged (**a**, n = 14,342), higher (**b**, n = 10,313), and lower (**c**, n = 7,411).

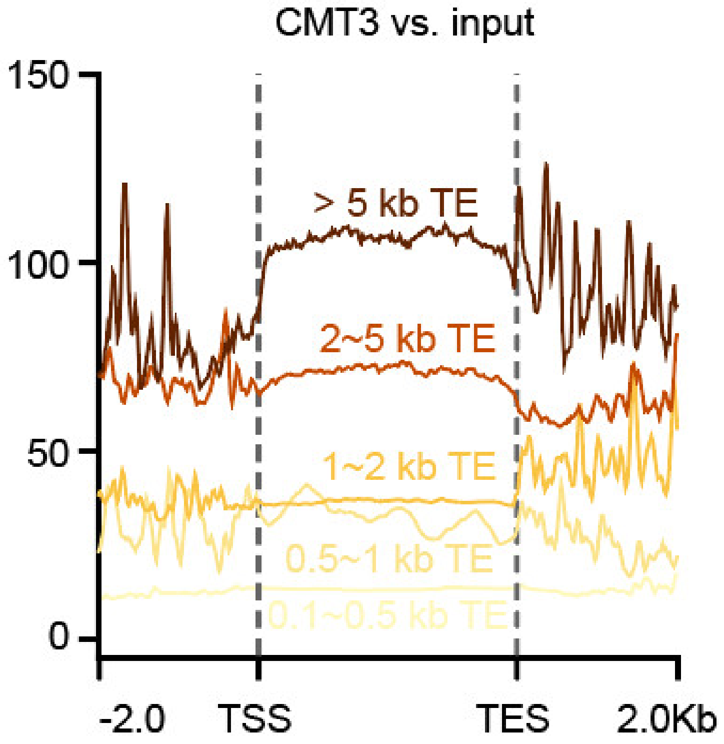


**Supplementary Fig. 3 CMT3 binding over TEs.** Metaplot showing CMT3 ChIP-seq signal (log2 FLAG/input) over transposons grouped by lengths. TEs are divided into five groups: 0.1 to 0.5 kb TEs (n=12,018), 0.5 to 1 kb TEs (n=5,856), 1 to 2 kb TEs (n=3,471), 2 to 5 kb TEs (n=2,225), and > 5 kb TEs (n=1,022).

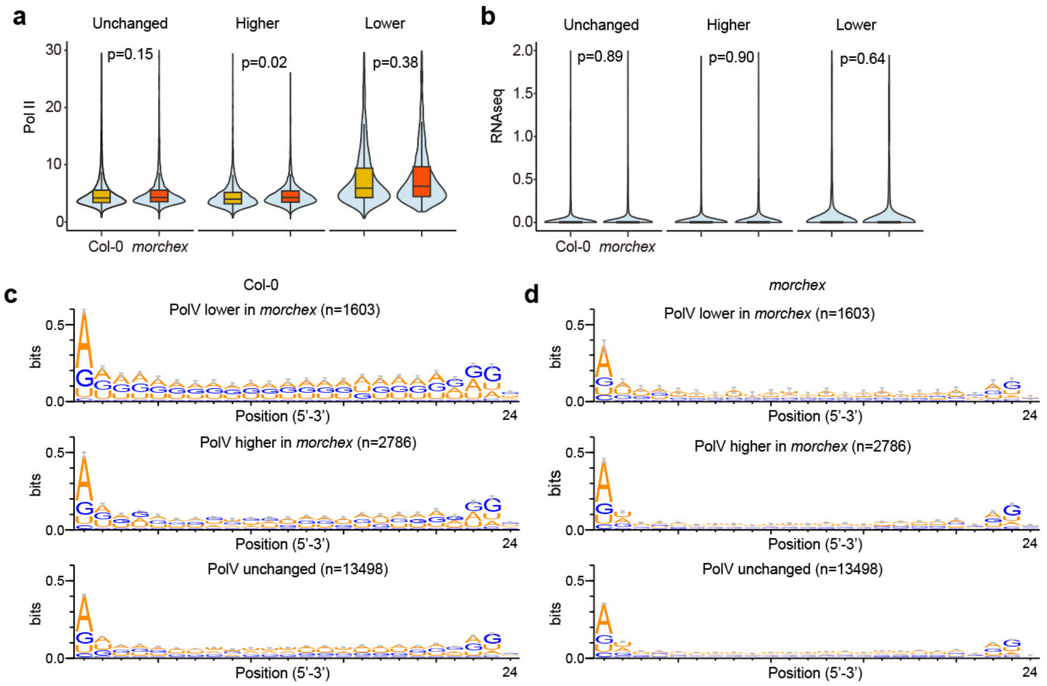


**Supplementary Fig. 4 a.** Violin plot illustrating Pol II binding level over Pol V lower, Pol V higher, and Pol V unchanged peaks. **b.** Violin plot displaying poly-A RNA transcription levels over Pol V lower, Pol V higher, and Pol V unchanged peaks. **f-g.** Nucleotide bias of small RNA of Pol V lower, Pol V higher, and Pol V unchanged peaks in Col-0 and *morchex* mutants.

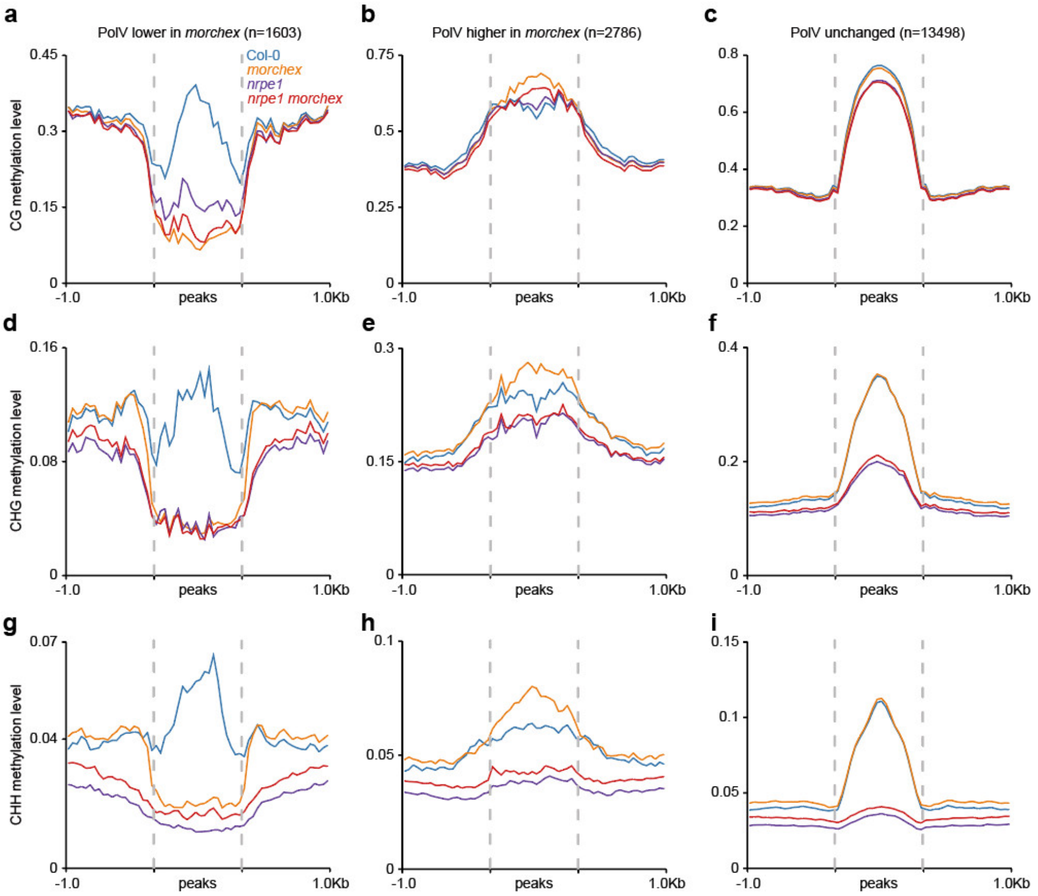


**Supplementary Fig. 5 DNA methylation dynamics associated with Pol V redistribution in *morchex* mutant.** CG methylation level over Pol V lower (**a**), Pol V higher (**b**), and Pol V unchanged peaks (**c**) in WT, *morchex*, *npre1*, and *nrpe1 morchex* mutants. CHG methylation level over Pol V lower (**d**), Pol V higher (**e**), and Pol V unchanged (**f**) peaks in WT, *morchex*, *npre1*, and *nrpe1 morchex* mutants. CHH methylation level over Pol V lower (**g**), Pol V higher (**h**), and Pol V unchanged (**i**) peaks in WT, *morchex*, *npre1*, and *nrpe1 morchex* mutants.
